# Sex-differences in prostaglandin signaling: a semi-systematic review and characterization of PTGDS expression in human sensory neurons

**DOI:** 10.1101/2022.11.25.517978

**Authors:** Breanna Q. Shen, Ishwarya Sankaranarayanan, Theodore J. Price, Diana Tavares-Ferreira

**Affiliations:** University of Texas at Dallas, Department of Neuroscience and Center for Advanced Pain Studies; Richardson, TX, USA

## Abstract

There is increasing evidence of sex differences in underlying mechanisms causing pain in preclinical models, and in clinical populations. There are also important disconnects between clinical pain populations and the way preclinical pain studies are conducted. For instance, osteoarthritis pain more frequently affects women but most preclinical studies have been conducted using males in animal models. The most widely used painkillers, nonsteroidal anti-inflammatory drugs (NSAIDs), act on the prostaglandin pathway by inhibiting cyclooxygenase (COX) enzymes. The purpose of this study was to analyze the preclinical and clinical literature on the role of prostaglandins and COX in inflammation and pain. We aimed to specifically identify studies that used both sexes and investigate whether any sex-differences in the action of prostaglandins and COX inhibition had been reported, either in clinical or preclinical studies. We conducted a PubMed search and identified 369 preclinical studies and 100 clinical studies that matched our inclusion/exclusion criteria. Our analysis shows that only 17% of preclinical studies on prostaglandins used both sexes and, out of those, only 19% analyzed or reported data in a sex-aware fashion. In contrast, 79% of the clinical studies analyzed used both sexes. However, only 6% of those reported data in a sex-aware fashion. Interestingly, 14 out of 15 preclinical studies and 5 out of 6 clinical studies that analyzed data in a sex-aware fashion have identified sex-differences. This builds on the increasing evidence of sex-differences in prostaglandin signaling and the importance of sex-awareness in data analysis. The preclinical literature identifies a sex difference in prostaglandin D_2_ synthase (PTGDS) expression where it is higher in female than in male rodents in the nervous system. We experimentally validated that PTGDS expression is higher in female human dorsal root ganglia (DRG) neurons recovered from organ donors. Our semi-systematic literature review reveals a need for continued inclusivity of both male and female animals in prostaglandins studies and sex-aware analysis in data analysis in preclinical and clinical studies. Our finding of sex-differences in neuronal PTGDS expression in humans exemplifies the need for a more comprehensive understanding of how the prostaglandin system functions in the DRG in rodents and humans.

## INTRODUCTION

Differences in the incidence and severity of many pain disorders between men and women have been reported. The prevalence of fibromyalgia [2] migraine [60], and osteoarthritis pain [61] is greater among women than men. Women are also reported to have more severe postoperative pain [16]. The chronic pain patient population is largely female, older, and genetically heterogeneous, while preclinical pain studies have been primarily conducted in young, male mice or rats of limited, and usually inbred strain [28]. The past decade has seen an explosion of preclinical pain research demonstrating fundamental sex differences in mechanisms causing chronic pain in mouse and rat models [29; 45]. Some female-specific pain mechanisms are now emerging in the preclinical literature, such as the more prominent effects of calcitonin gene-related peptide and prolactin in promoting pain in female rodents [4; 5; 37; 38]. Similar differences are now emerging at the molecular level in the human dorsal root ganglion (DRG) [42; 43; 56] suggesting that sex differences in basic pain mechanisms may contribute to differential efficacy of pain therapeutics in men and women. We recently identified a sex difference in DRG neuron expression of a prostaglandin synthesizing enzyme, PTGDS, that led to differences in behavioral outcomes in response to prostaglandins and PTGDS inhibitors in male and female mice [55]. This finding prompted us to look more carefully at the clinical and preclinical research on prostaglandins and their receptors with the hypothesis that the very commonly used drugs that target this pathway may have differential efficacy in men versus women.

Prostaglandins (PGs) are lipid-derived signaling molecules that play an important role in pain and inflammation. PGs are produced from plasma membrane-derived arachidonic acid, and their conversion is dependent on the action of cyclooxygenase (COX) enzymes (**Figure 1**). The four major prostaglandins, prostaglandin E2 (PGE_2_), prostaglandin D2 (PGD_2_), prostacyclin (PGI_2_), and prostaglandin F_2a_ (PGF_2a_) act on G-protein coupled receptors to regulate intracellular signaling pathways [44]. PGE_2_ acts on the E prostanoid receptors EP1, EP2, EP3, and EP4. As Gs-coupled receptors, EP2 and EP4 activate adenylyl cyclase to produce cyclic adenosine monophosphate (cAMP) from adenosine triphosphate (ATP); as a second messenger, cAMP phosphorylates protein kinase A (PKA), which in turn can phosphorylate intracellular target proteins [44]. EP3 is a G_i_- and G_12_-coupled, lowering cAMP, increasing intracellular calcium and the activity of rho GTPases [44] and this receptor has been linked to anti-nociceptive actions of PGE_2_ [31]. EP1 is a G_q_-coupled receptor, whose activation causes phospholipase C to catalyze conversion of phosphatidylinositol biphosphate (PIP_2_) into inositol triphosphate (IP3) and diacylglycerol (DAG). All 4 EP receptor genes are expressed in mouse [70] and human [56] DRG neurons where their expression varies depending on the subtype of receptor and the type of sensory neuron. EP1, 2 and 4 receptors are known to sensitize ion channels like the TRPV1 receptor and voltage gated sodium channels that regulate the excitability of sensory neurons [10; 14].

**Figure 1.**
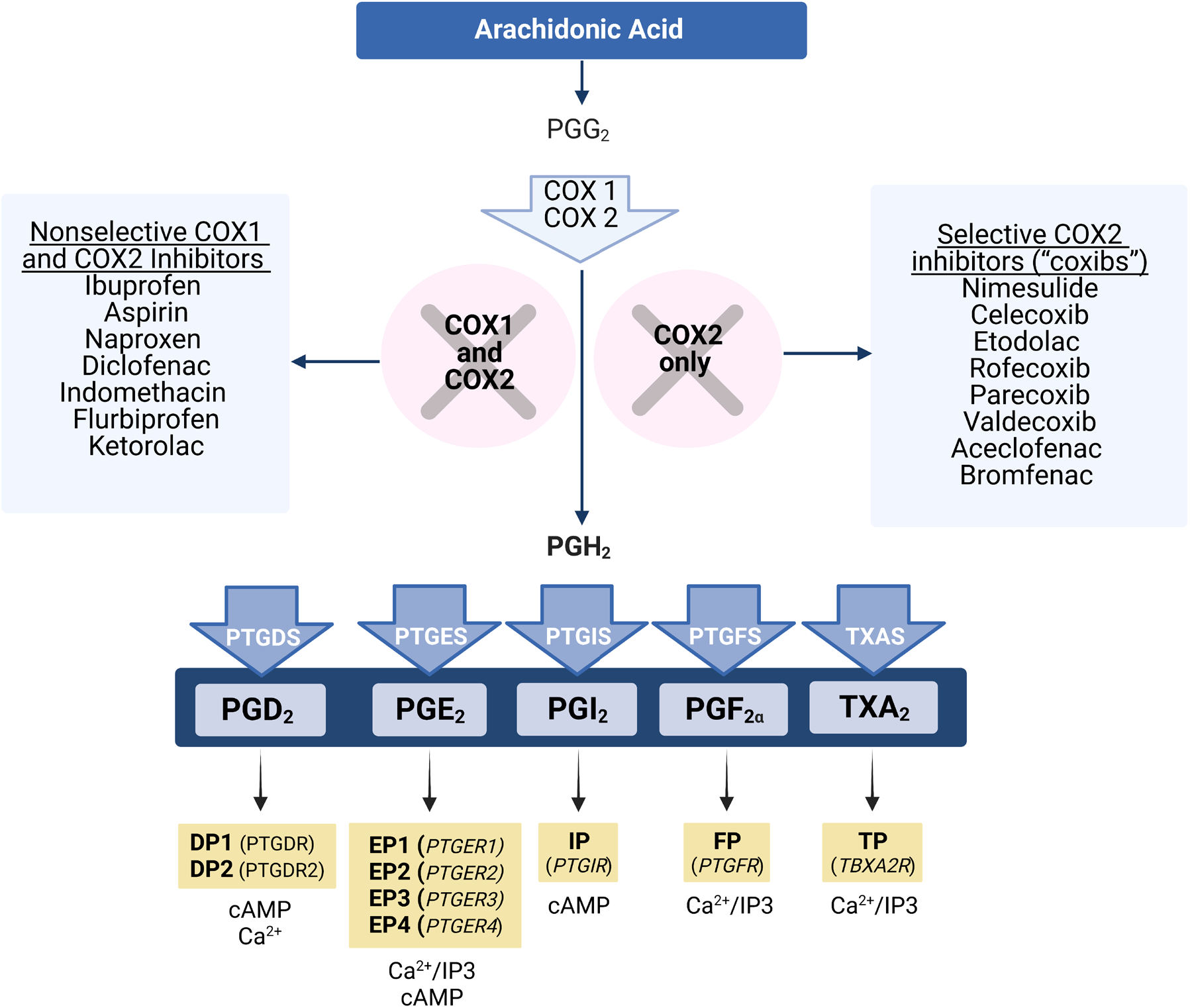
Prostaglandins biosynthesis pathway. Cyclooxygenases (COX) metabolize arachidonic acid first to prostaglandin G_2_ (PGG_2_) and then to prostaglandin H_2_ (PGH_2_). COX1 and 2 are targeted by non-steroidal anti-inflammatory drugs (NSAIDs), which block the synthesis of prostaglandins. PGH_2_ is converted to prostaglandin D_2_ (PGD_2_), prostaglandin E_2_ (PGE_2_), prostaglandin I_2_ (PGI_2_), prostaglandin F_2α_ (PGF_2α_) and thromboxane A_2_ (TXA_2_) by respective synthases. Each prostaglandin binds to specific receptors, which are members of the G protein-coupled receptor (GCPR) superfamily of seven transmembrane proteins, activating different downstream signaling pathways. cAMP: cyclic adenosine monophosphate; IP3: inositol triphosphate.

PGD_2_ acts on two GPCRs, DP1, a G_s_-coupled receptor that increases cAMP concentration, and DP2, a G_i_-coupled receptor also known as chemoattractant receptor-homologous molecule expressed on T helper 2 cells (CRTH_2_) that decreases cAMP concentration and increases calcium intracellularly [44]. While some studies have shown that PGD_2_ is pro-nociceptive [15], other studies have demonstrated that PGD_2_ has anti-nociceptive effects [26]. PGI_2_ exerts pro-inflammatory effects through the prostacyclin receptor (IP), a G_s_-coupled receptor that increases cAMP concentration [44]. IP mediates both peripheral nociception and spinal transmission of nociceptive information [44]. In dorsal root ganglia (DRG) and spinal dorsal horn neurons, IP receptor activation causes increased neuronal excitability [7]. In addition, IP has an important role inducing inflammation, causing vasodilation and edema [44]. PGF_2a_ activates the FP_A_ and FP_B_ receptors, which are G_q_-coupled and increase concentrations of IP3, DAG, calcium, and rho activity [44]. PGF_2a_ has pro-inflammatory effects in arthritic conditions and can induce inflammation, but the mechanisms are unclear [44]. Finally, an additional prostanoid, thromboxane A2 acts on TP_a_ and TP_b_ GPCRs to increase IP3, DAG, calcium, and activity of RhoGEF in cells [44].

Nonsteroidal anti-inflammatory drugs (NSAIDs) reduce inflammation and its symptoms through either nonselective inhibition of both COX isoforms or selective inhibition of COX2. COX inhibition then reduces the production of PGs. COX-1 is ubiquitously expressed and plays a key role in maintenance of mucosal tissues, and COX-2 is upregulated by injury and contributes to inflammation [44]. NSAIDs were first used in the form of myrtle and willow tree bark by the Egyptian civilization around 1500 BC for their analgesic and antipyretic functions, and salacin was first isolated from the willow bark in 1828 [62]. Salicylic acid (aspirin) was subsequently synthesized and marketed, followed by ibuprofen and around 50 other NSAIDs. Globally, NSAIDs are some of the most widely used and prescribed drugs [62]. While aspirin is a noncompetitive COX inhibitor, non-aspirin NSAIDs competitively inhibit active sites on COX enzymes [62]. In an attempt to eliminate the gastrointestinal complications of COX-1 inhibition, COX-2 selective inhibitors, also known as coxibs, were developed in the 1990s [22].

The primary goal of our semi-systematic review was to first identify if males, females or both sexes have been used in studies investigating PGs in pain and inflammation. Second, we aimed to identify any reports of sex-differences in PG signaling in the preclinical literature (studies using non-human subjects) and at the clinical level. Our semi-systematic review found that there is a large bias towards using male animals in prostaglandin preclinical studies. While most clinical studies include both male and female subjects, only a small proportion of studies reported data in a sex-aware fashion. To provide further evidence of the need to look at the impact of sex differences in studies on prostaglandins and pain, we characterized the expression of PTGDS, an enzyme that converts PGH_2_ to PGD_2_, in human DRG neurons. We found that PTGDS protein expression is higher in DRG neurons recovered from female organ donors compared to male organ donors. We conclude that potential sex differences in the action of prostaglandins should be studied more carefully in preclinical and clinical studies moving forward.

## MATERIALS AND METHODS

### Preclinical studies

#### Pubmed search

A total of 1,825 articles were retrieved from PubMed on November 20, 2020 using the following keywords: “prostaglandins, inflammation, and pain”. 369 papers were included for analysis (**Figure 2A**).

**Figure 2.**
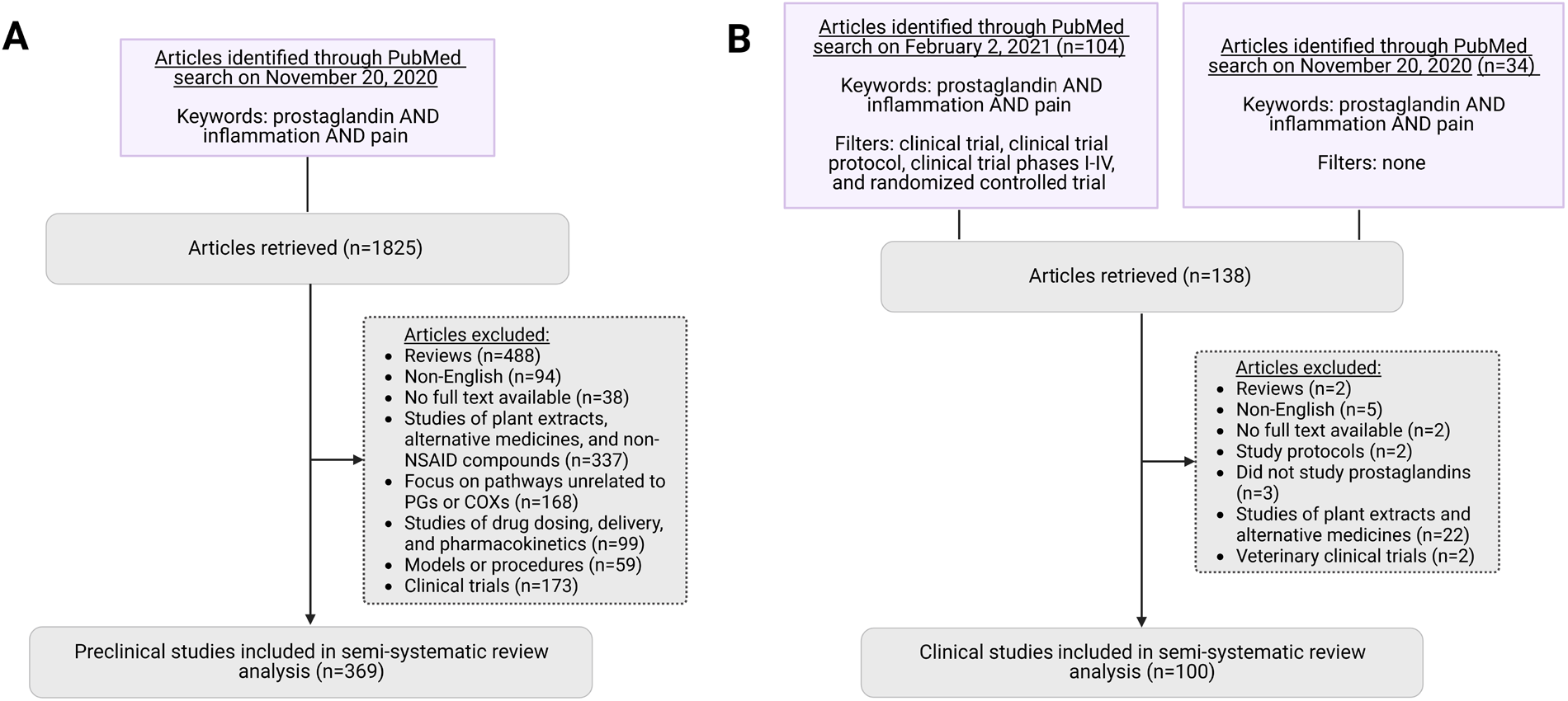
Semi-systematic review of articles retrieved on PubMed. A) Workflow of pre-clinical studies identified through PubMed. B) Workflow of clinical studies identified through PubMed.

#### Inclusion criteria

Articles of preclinical studies were found using the keywords mentioned above. Only papers that involved animal models and/or the use of human tissues were included. Only articles with PMID were included. A total of 369 papers were included.

#### Exclusion criteria

Papers were excluded if they were reviews (n=488), not written in English (n=94), had no full text available (n=38), studied plant extracts, alternative medicines, and other non-NSAID compounds (n=337), focused on pathways unrelated to PGs or COXs (n=168), studied drug dosing, delivery, or pharmacokinetics only (n=99), presented models or procedures only (n=59), or were clinical trials (n=173).

#### Data collection

A semi-systematic analysis was conducted for papers that analyzed data in a sex-aware fashion. Information such as the specific PGs, COX enzymes, NSAIDs, and receptors investigated in each study were recorded.

#### Data analysis

The number of studies conducted each year with males, females, or both was recorded. For papers that included both male and female subjects, we also noted whether data was analyzed in a sex-aware fashion. Finally, we examined any sex differences in PG or COX inhibitor actions (**Suppl. Files 1 and 2**).

### Clinical Trials

#### Pubmed search

A total of 104 papers on prostaglandins, inflammation, and pain were retrieved from PubMed on Feb 2, 2021. Papers were clinical trials, clinical trial protocols, clinical trial phases I-IV, and randomized controlled trials (**Figure 2B**). An additional 34 papers on the same topic were retrieved from a PubMed search on November 20, 2020, without the above filters.

#### Inclusion criteria

Clinical trial articles were included upon mention of all three search criteria, prostaglandins, inflammation, and pain. Included studies must also involve human subjects. Only articles with PMID were included. A total of 100 papers were included.

#### Exclusion criteria

Papers were excluded if they were reviews (n=2), not written in English (n=5), had no full text available (n=2), described study protocols only (n=2), did not study prostaglandins (n=3), studied plant extracts or alternative medicine treatments (n=22), or were veterinary clinical trials (n=2).

#### Data collection

A semi-systematic analysis was conducted for all articles regardless of sex-aware analysis. We recorded the specific PGs, COX enzymes, NSAIDs, and receptors investigated. We also identified the purpose, experimental design, model, sex, number, group design, doses, injection type, measurements and tissues collected.

The papers analyzed include either clinical pain models (studies involving patients diagnosed with a pain condition, n=74) or experimental pain conditions (n=26). Experimental pain conditions include eccentric exercise-induced muscle pain (n=4), experimentally induced skin hyperalgesia (n=20), and experimental models of sleep deprivation (n=2). All other studies involve clinical pain conditions, such as surgeries, joint pain, and painful dental conditions.

#### Data analysis

We categorized papers by clinical model and compared the main findings. Any discrepancies and/or agreements between studies were investigated in terms of divergent experimental methodology. We also recorded the number of clinical trials conducted each year with males, females, or both. Finally, we examined any sex differences in PG or COX inhibitor action (**Suppl. Files 3 and 4)**.

### Immunohistochemistry

Human dorsal root ganglia (DRG) from the L4 and L5 levels were recovered from organ donors, frozen immediately on crushed dry ice, and stored in a -80 C freezer as previously described [49]. The DRGs were embedded in OCT by gradually adding layers of OCT on the tissue, which was kept frozen over dry ice. Tissues were sectioned in the cryostat at 20 µm and adhered onto SuperFrost Plus charged slides (Thermo Fisher Scientific). Slides were kept in the -20 C cryostat chamber for 15 minutes following completion of sectioning. The slides were then immediately fixed in ice-cold formalin (10%) for 1 minute followed by dehydration in 50% ethanol (1 minute), 70% ethanol (1 minute), and 100% ethanol (2 minutes) at room temperature. The slides were briefly air dried. A hydrophobic pen (ImmEdge PAP Pen; Vector Labs) was used to draw boundaries around each tissue section, and boundaries were allowed to air dry.

Slides were incubated with blocking buffer (10% Normal Goat Serum, Atlanta Biologicals, Cat #S13150h, 0.3% Triton X-100 in 0.1 M PB) for 1 hour at room temperature. Sections were then incubated overnight with a primary antibody cocktail. Following primary antibody incubation, sections were washed with 0.1 M phosphate buffer and incubated with Alexa Fluor secondary antibodies (Fisher Scientific/Invitrogen, dilutions 1:1000) for 1 hour at room temperature. Sections were washed in 0.1 M phosphate buffer. To remove lipofuscin signal, Trublack (1:20 in 70% ethanol; Biotium #23007) was pipetted to cover each section for 1 minute before being rinsed off. Finally, slides were air dried and cover slipped with Prolong Gold Antifade reagent (Fisher Scientific; P36930). The PTGDS antibody (ab18214, dilution 1:100) was obtained from Abcam and the peripherin antibody (P5117, dilution 1:500) was obtained from Sigma-Aldrich. The DP1 antibody (101640, dilution 1:200) was obtained from Cayman Chemicals and the SOX10 antibody (ab216020, dilution 1:40) was obtained from Abcam.

### Image acquisition and PTGDS quantification

DRG sections with PTGDS staining (n=12) were imaged on an Olympus VS120 Virtual Slide Microscope and Olympus FluoView 1200 confocal microscope, using the same settings for all images. Images were analyzed using Olympus CellSens software. The mean gray intensity of all neurons in a full DRG section from each donor was quantified (on average, we quantified 619 neurons per DRG section; the lowest number of neurons per DRG was 268). DRG sections with DP1 staining were imaged on an Olympus FV3000RS Confocal Laser Scanning Microscope.

### Statistics

Statistical analysis for PTGDS quantification was done in GraphPad Prism 9.3.1. The mean gray intensity values were normalized by the area of the neurons. Single comparisons were performed on all neurons using Student’s t test with Welch’s correction (which accounts for the inequal variances per group). For statistical analysis, data from the 61-year-old female (donor #12) was excluded (post-menopause). In total 6813 neurons were analyzed (2920 from 5 female DRG samples and 3893 from 6 male DRG samples). Statistical results can be found in figure 4 legend.

Data visualization for figure 3 was done in Python (version 3.8.5 with Anaconda distribution). Figures 1 and 2 were generated using Biorender (BioRender.com).

**Figure 3.**
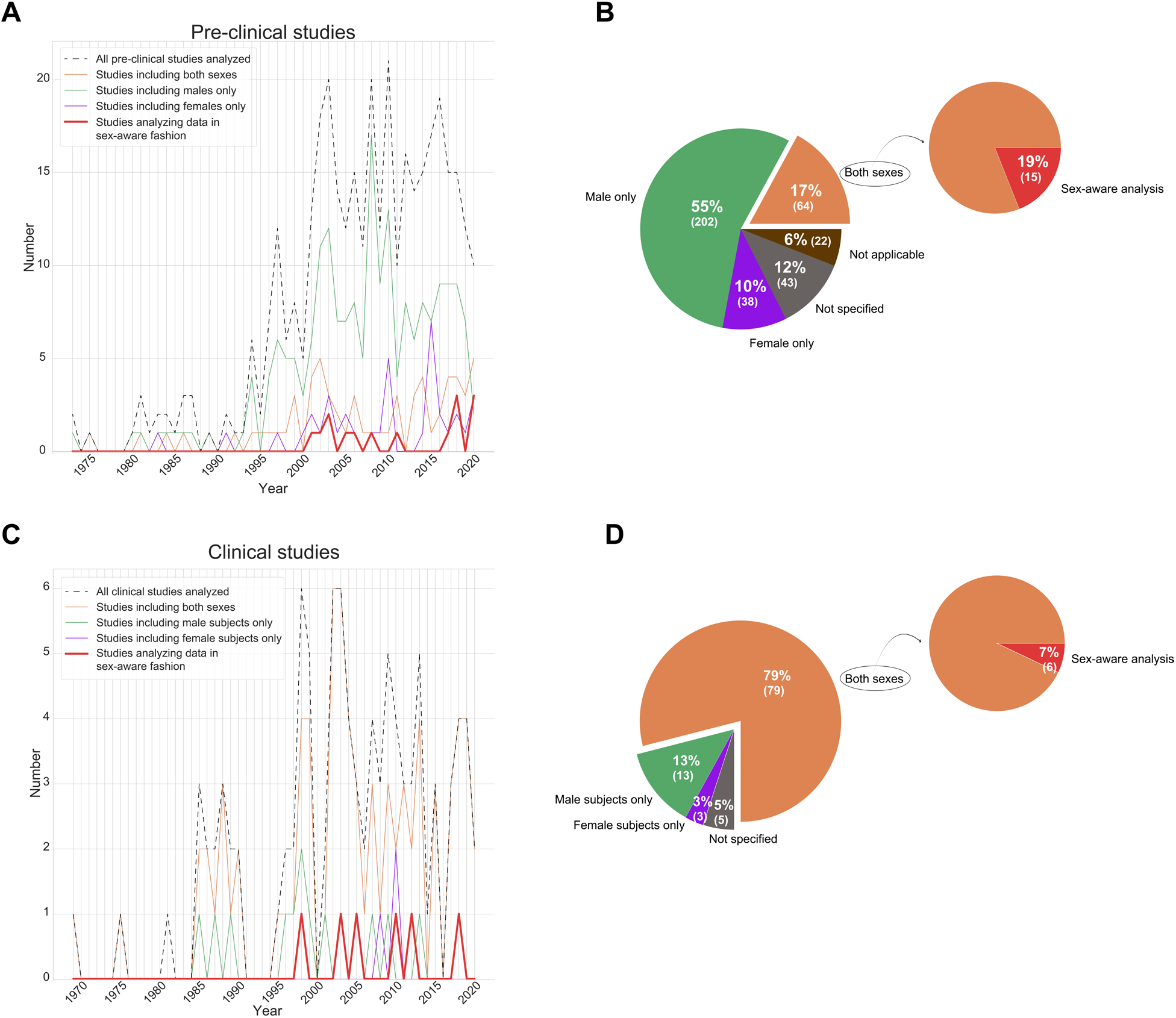
Most prostaglandin studies have not been analyzed in a sex-aware fashion. **A)** Line plot showing the inclusion of male, female or both sexes in pre-clinical studies from 1973 to 2020 and the number of studies that have been analyzed in a sex-aware fashion (red line). **B)** Pie charts showing the overall percentage of pre-clinical studies that include male, female or both sexes from 1973 to 2020. **C)** Line plot showing the inclusion of male, female or both sexes in clinical studies from 1969 to 2020, including the studies that have analyzed the data in a sex-aware fashion (red line). **D)** Pie charts showing the overall percentage of clinical studies that include male, female or both sexes from 1969 to 2020. Note: studies where the sex of animals/subjects is not clear or specified are not shown in the line plots, please refer to supplementary files 1-4.

## RESULTS

We conducted a semi-systematic review of the literature on prostaglandins (PGs), inflammation and pain. Papers were categorized as preclinical studies, which included animal models and/or human tissues, and as clinical studies, which included human subjects in clinical or experimental settings.

### Preclinical Studies

Details on the preclinical studies included in our semi-systematic review can be found in **supplementary files 1 and 2**. Here we summarize the articles that report results in a sex-aware fashion.

In several studies, male and female animals demonstrated differences in response to noxious substance exposure related to prostaglandins. In a model of IL-1b induced temporomandibular joint (TMJ) inflammation, female rats showed greater sensitivity in the TMJ region than male rats before and after implantation of a cannula. Female rats also had a lower head withdrawal threshold after intratrigeminal ganglionic IL-1b injection but this difference was not statistically significant. IL-1b upregulates COX2 mRNA and protein, resulting in upregulation of PGE2, increased EP2 activation, and increased Nav1.7 expression. No significant differences were observed in expression fold change for Nav1.7 and COX2 mRNA and protein between males and females [71]. While there is inconclusive evidence on sex differences in the endogenous production of PGE_2_ in response to introduction of IL-1b, application of exogenous PGE_2_ as a component of a combination of inflammatory mediators may cause sex-dependent sensitization of neurons. In a different study, the proportion of dural afferents sensitized by an inflammatory mediator solution of bradykinin, histamine, and PGE_2_ was significantly greater in female rats than male rats. The investigators suggest this may be caused by different changes in TTX-R sodium current of males and females [46]. Since three different inflammatory mediators were used in the study, it is not possible to know whether sex-differences are solely caused by PGE_2_.

Sex differences in the translatomes (RNAs bound to ribosomes) and transcriptomes of male and female mice DRG have been reported. In the prostaglandin pathway, prostaglandin D2 synthase (*Ptgds*) was found to be upregulated in female DRG neurons. Female mice also exhibited higher levels of PTGDS protein and PGD_2_ [55]. PTGDS blockade produced more intense grimacing in male compared to female mice, suggesting that endogenous PGD_2_ might reduce nociception in the absence of injury. In the same study, it was also observed that female mice displayed more mechanical allodynia and grimacing after PGE_2_ injection than male mice. This study indicates that there is a complex balance between different prostaglandins in regulating nociceptive behavior in mice.

In studies investigating COX enzymes, female COX1 knockout mice and female COX2 knockout mice showed reduced edema and joint destruction in a model of Freund’s adjuvant-induced arthritis compared to male mice. Female COX1 knockout mice also showed reduced contralateral allodynia compared with males [12]. Similarly, another study demonstrated a reduced writhing response to acetic acid in COX2 deficient female mice. However, the anti-nociceptive effect was not seen in COX2 deficient male mice [40]. On the other hand, NSAIDs that antagonize COX action appear to be less effective in females. The COX2 inhibitor celecoxib generated shorter and less prominent effects in female than male mice, when administered after Complete Freund’s Adjuvant injection [24]. COX inhibitors may also have different anti-inflammatory effects in males and females. In rats, indomethacin, a COX inhibitor, administered after experimental surgery decreased the elevated interleukin-6 levels in males only [36]. These seemingly conflicting results present an interesting paradox: while reduction/ablation of COX2 expression through genetic manipulation may be more effective in preventing nociceptive effects in females, reduction of COX2 through pharmacological inhibitors, such as NSAIDs, may be more effective in males. Several factors may contribute to this effect, including sex-differences in expression of genes related to drug metabolism and drug transporter expression [17]. In fact, the activity level of cytochrome p450 34A, an enzyme involved in drug metabolism, is higher in females [52]. This could be consistent with lower COX-inhibitor efficacy in females, but the effects may also be mediated by differences in signaling at the level of the DRG nociceptor, as suggested by other studies described above.

### Clinical Studies

Clinical trials were subdivided in two ways. First, studies were categorized by how the pain conditions occurred: experimentally in the laboratory (n=26) or clinically due to a disease process and/or procedural intervention (n=74). Second, they were categorized by clinical model of disease and compared within these groups. Clinical trials were grouped by clinical model as follows: oral health (n=10), surgical extraction of third molars (n=14), hip and knee surgeries (n=14), general surgeries (n=13), tendon and joint disorders (n=7), muscle pain (n=7), experimentally induced skin hyperalgesia (n=20), vascular system diseases (n=5), experimental models of sleep deprivation (n=2), headache (n=2), and general studies of NSAIDs (n=6). This grouping scheme allowed us to compare and contrast conclusions derived from relatively uniform settings. More details on the included clinical studies can be found in **Supplementary Files 3 and 4**.

Most clinical studies using the oral health model investigated whether levels of PGE_2_ can be correlated with severity of gum disease and patient-reported pain. Other studies focused on the efficacy of COX inhibitors. However, no studies reported data in a sex-aware fashion. One study found that there is variability in response to COX inhibitors. In third molar extraction surgery, some participants are partial responders to ibuprofen (4 men, 5 women), and others are complete responders (7 men, 3 women) [57]. While the ratio of men to women in the complete responder group is greater than in the partial responder group to ibuprofen, the study did not analyze the difference in complete and partial responders in terms of potential sex differences, discussing only differences in levels of PG metabolites, cytokines, peripheral blood mononuclear cells gene expression between the two groups (**Suppl. File 3**) [57].

The analgesic and anti-inflammatory effects of various coxibs and nonselective COX inhibitors were compared in patients undergoing hip and knee treatment, e.g. total knee arthroplasty and hip replacement due to osteoarthritis. Only one study on knee arthroscopy observed that the women were at greater risk of developing moderate or severe pain [51]. Other studies involving patients undergoing hip and knee procedures included both men and women but did not analyze data in a sex-aware fashion or report sex-differences.

Experimental skin hyperalgesia was induced in several studies by injections of PGE_2_ or low pH solutions into the muscle and skin, potentiating nociceptor activation [30]. In a model of experimentally induced sunburn (UVB-induced erythema), ibuprofen (COX inhibitor) increased heat pain threshold and heat pain tolerance overall [53]. However, the authors noted that men were more responsive to ibuprofen compared to women, which suggests that the anti-inflammatory effect is greater in men. Similarly, in a model of electrically induced pain, only men showed response to ibuprofen treatment and had higher pain tolerance [63] [9].

Two studies on headache investigated whether headache is induced by inflammatory mediators [1; 3]. The authors did not report data in a sex-aware fashion.

### Sex-awareness in subject selection and data analysis

#### Approximately 17 % of preclinical studies examined showed sex-awareness in subject selection and, out of those, 19% showed sex-awareness in data analysis

Preclinical studies analyzed in this semi-systematic review were published in years ranging from 1973 to November 2020. Among the 369 total studies analyzed, 202 studies involved the use of male animals only, 38 studies involved the use of female animals only, and 64 studies included the use of both male and female animals (**Figure 3A, B; Suppl. File 3**). Several of these studies included both males and females, but in different experiments within the study, resulting in no direct comparisons. Additionally, 43 studies contained unclear information on the sex of the experimental animals, and 22 studies involved experimental setups where the sex information was not applicable, such as cell cultures (**Suppl. File 3**).

Out of the 369 total studies analyzed, only 64 studies were conducted using both male and female experimental animals. Out of the 64 studies that included male and female subjects, only 15 studies analyzed data in a sex-aware fashion, regardless of whether they found significant sex differences or not. While 14 of these 15 studies included comments comparing prostaglandin-related action in males and females, one study presented data from each human tissue donor separately without further analyses [8]. We note that although studies in our analysis ranged from 1973-2020, all 15 studies that analyzed data in a sex-aware fashion were published after 2000 (one in 2001, one in 2002, two in 2003, one in 2005, one in 2006, one in 2008, one in 2011, one in 2017, three in 2018, and three before our cutoff date in 2020). This suggests that the relevance of sex-aware analyses is becoming more widely recognized. In addition, 12 out of the 15 studies that showed sex-awareness in data analysis included animals only, and the remaining 3 out of these 15 studies included human tissues only. However, a much larger proportion of the studies that included both male and female subjects made at least some use of human tissues, at 25 out of 64 studies. This suggests that inclusion of human tissues from both male and female donors does not necessarily translate into greater sex-awareness in data analysis. An alternative interpretation is that there were no sex differences but the authors did not report their negative findings on the lack of sex difference. The 15 preclinical studies that analyzed data in a sex-aware fashion found many differences between male and female animals in the prostaglandin pathways, behavior in response to noxious stimuli, etc. which we described in the first part of our results.

#### 79% of clinical studies examined showed sex-awareness in subject selection and, out of those, approximately 7% showed sex-awareness in data analysis

Contrary to preclinical studies, most clinical trials include both male and female participants (**Figure 3C, D; Suppl. File 4**). However, few studies reported results separately for men and women. It is unclear whether the data from men and women were merged because no sex differences were found or if potential sex differences were not assessed at all. Some studies included subjects of only one biological sex. Similar to preclinical studies, these single-sex studies in the experimental pain category showed a bias towards using male subjects (six studies involved men only, and none involved women only). In clinical pain conditions, the 71 studies that included both male and female subjects showed highly variable proportions of male to female participants. Numbers range from 187 men: 9 women (95.4% male) [19] to an equal split (10 men: 10 women) [13] to 3 men: 25 women [32]. Additionally, only one study using a clinical pain model analyzed data in a sex-aware fashion. This single study noted that female biological sex was a risk factor for moderate/severe postoperative pain after minor arthroscopic knee surgery [51]. Another study focusing on the investigation of NSAID use found that women report using more over the counter pain medications [67]. The remaining studies analyzed data from male and female participants together. In models of experimentally induced pain, the number of single-sex studies including male subjects only (n=13) and female subjects only (n=3) were also biased toward males only. Among the 4 experimental pain studies that analyzed data in a sex-aware fashion, there were some findings that appeared to be contradictory. In an eccentric exercise model, no sex differences were noted in changes in muscle function, muscle soreness, or histological data[39], even though serum creatine kinase pre-values and change after the first bout of exercise were lower in females [39]. On the other hand, a sunburn model showed that ibuprofen had a significantly greater effect on lowering skin temperature in men compared with women[53]. This finding in an inflammatory model agrees with similar conclusions made from two electrically-induced pain models. In these two electrical pain studies, ibuprofen had an analgesic effect only in males [63] [9].

### PTGDS is expressed in human DRG neurons and is enriched in neurons from female organ donors

Prostaglandin-D2 synthase (PTGDS) catalyzes the synthesis of prostaglandin D_2_ (PGD_2_), which is the most abundant prostaglandin in the brain [34; 48]. PGD_2_ controls nociception, sleep and temperature [15; 20; 23; 25; 27; 35; 47; 50; 58; 59; 64]. Previous research in our lab showed that PTGDS is enriched in female mouse DRG neurons [54]. We sought to investigate PTGDS expression in human DRG using immunohistochemistry (IHC). We found that PTGDS is colocalized with the neuronal marker peripherin and is expressed in human DRG neurons (**Figure 4A**). Next, we quantified PTGDS expression in human DRG from 12 organ donors (**Table 1**). We observed that PTGDS expression is higher in sensory neurons from female organ donors when compared to male organ donors (**Figure 4B, C**).

**Table 1.**
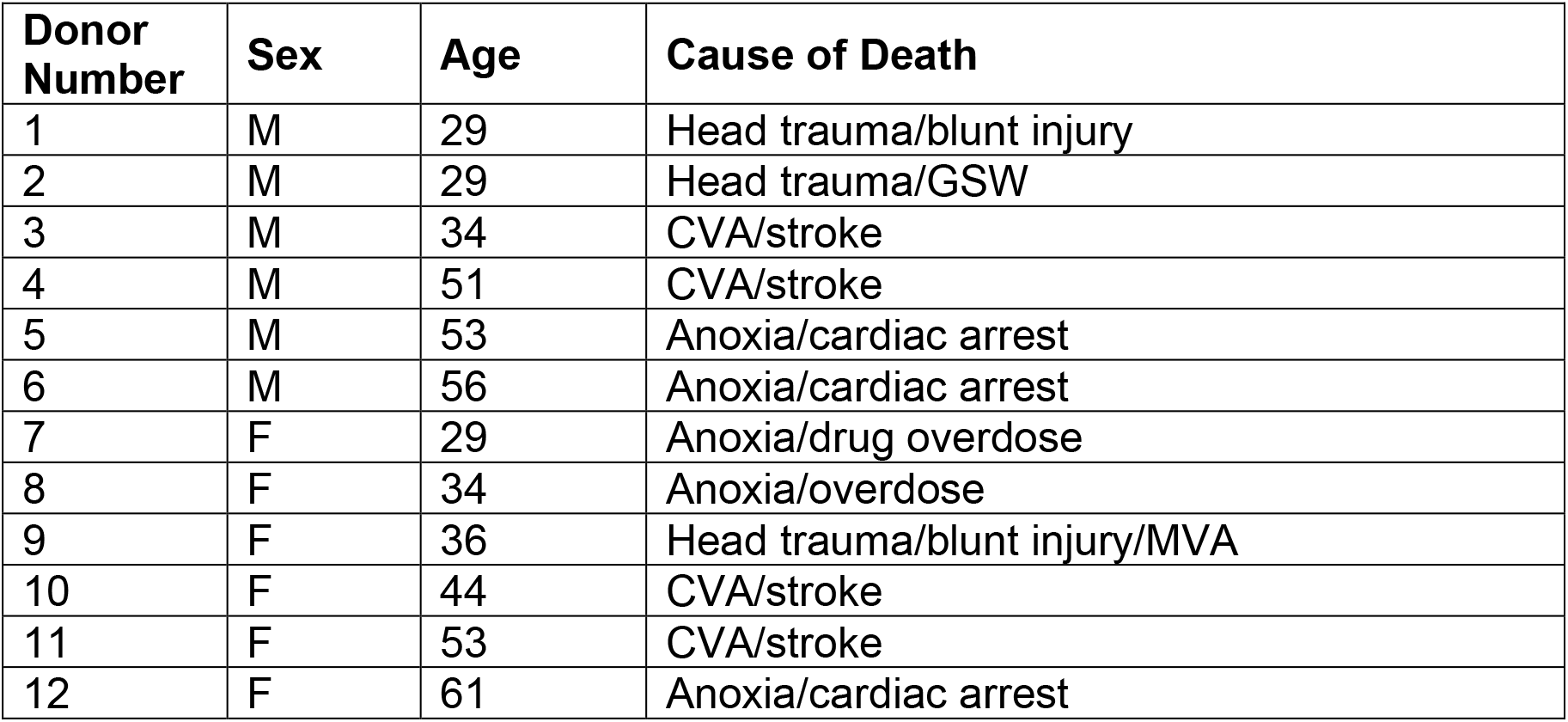
Donor information.

| **Donor Number** | **Sex** | **Age** | **Cause of Death** |
| --- | --- | --- | --- |
| 1 | M | 29 | Head trauma/blunt injury |
| 2 | M | 29 | Head trauma/GSW |
| 3 | M | 34 | CVA/stroke |
| 4 | M | 51 | CVA/stroke |
| 5 | M | 53 | Anoxia/cardiac arrest |
| 6 | M | 56 | Anoxia/cardiac arrest |
| 7 | F | 29 | Anoxia/drug overdose |
| 8 | F | 34 | Anoxia/overdose |
| 9 | F | 36 | Head trauma/blunt injury/MVA |
| 10 | F | 44 | CVA/stroke |
| 11 | F | 53 | CVA/stroke |
| 12 | F | 61 | Anoxia/cardiac arrest |

**Figure 4.**
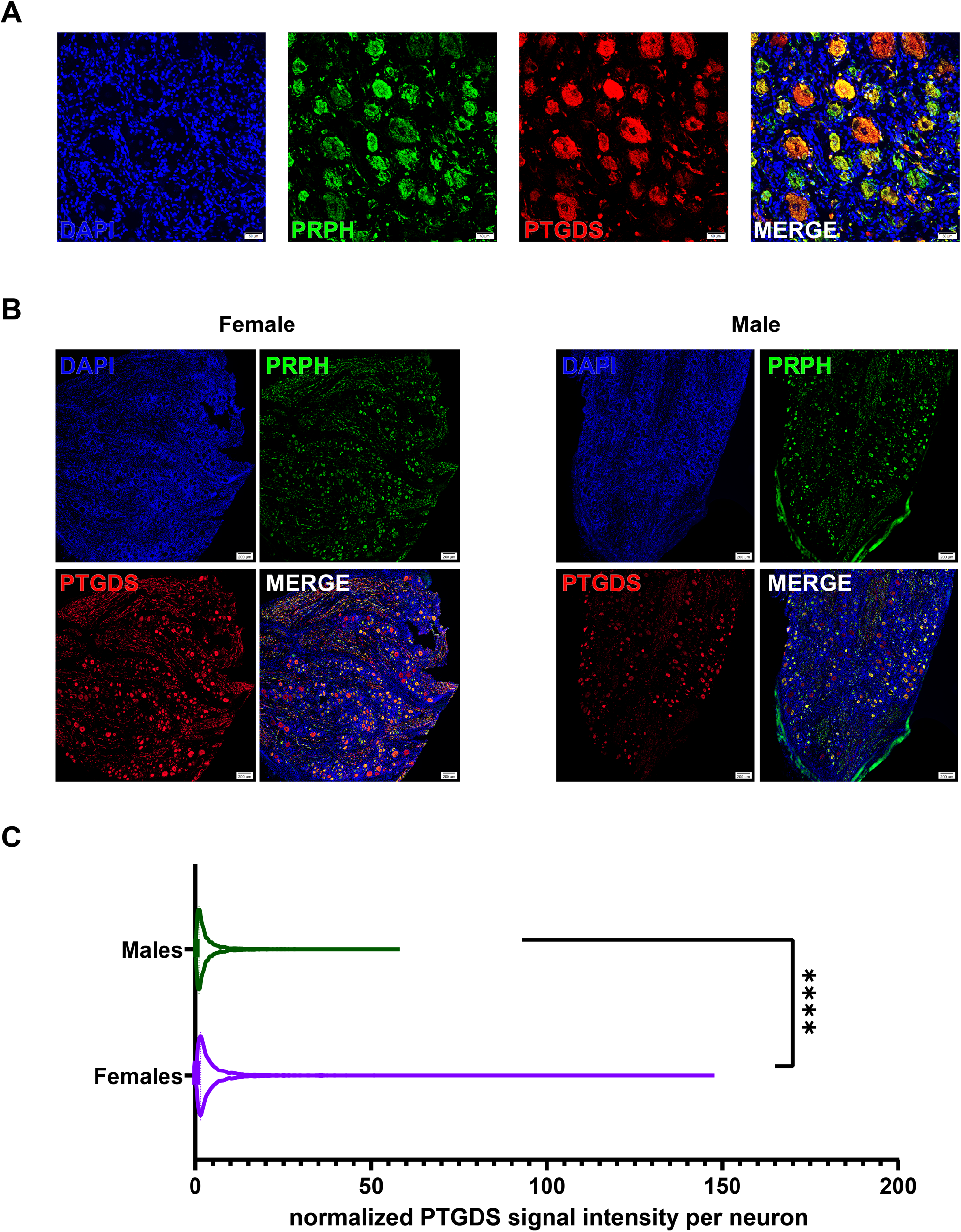
PTGDS is expressed in human DRG neurons. **A)** Confocal images show co-localization of PTGDS (red) and neuronal marker peripherin (PRPH, green). Cell nuclei are labelled with DAPI (blue). Scale bar = 50 µm. **B)** Expression of PTGDS in female (left) and male (right) DRG neurons. Scale bar = 200 µm. **C)** PTGDS has higher expression in female DRG neurons compared to male DRGs (Unpaired t test with Welch’s correction, t=11.08, df=4299, p-value<0.0001). **** p-value<0.0001. Signal intensity was normalized by the area of the neurons. Data from donor #12 was excluded (post-menopause, 61-year-old female). In total 6813 neurons were analyzed (2920 from 5 female DRG samples and 3893 from 6 male DRG samples).

### DP1 receptor expression in human DRG

PGD_2_ acts through two receptors, DP1 and DP2. Previous transcriptomic studies demonstrate that the gene that encodes DP2, *PTGDR2*, is scarcely detected in human DRG [41; 56; 65; 66]. Therefore, we decided to focus on characterizing the expression of DP1 in human DRG. Transcriptomic studies on human DRG do not suggest sex differences in *PTGDR1* expression, but the cell type expressing the gene is not clear from these studies [33; 43; 56]. Using IHC on both male and female DRG samples, our results suggest that DP1 is expressed mostly in glial cells surrounding neurons in human DRG, as it is highly colocalized with SOX10, a glial cell marker (**Figure 5**). These results suggest that PGD_2_ may be released by neurons and acts via DP1 that is expressed in glial cells surrounding neurons. Glial cells and their inter-communication with neurons have been previously reported to have a role in pain processing [11; 21; 68].

**Figure 5.**
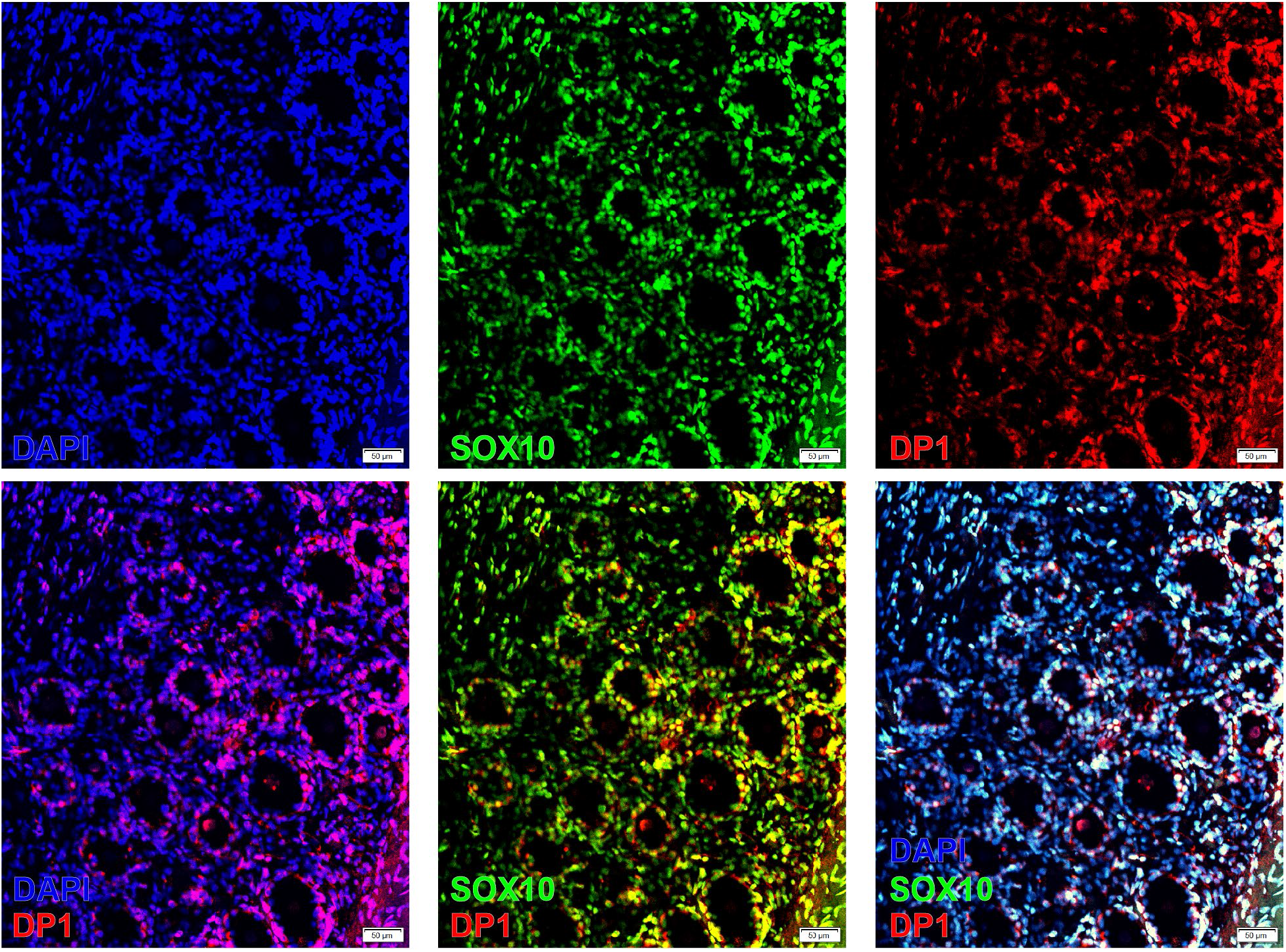
PGD2 receptor, DP1, is expressed in cells surrounding human DRG neurons. DP1 (red) is co-localized with SOX10+ (green) cells in human DRG. Cell nuclei are labelled with DAPI (blue). Scale bar = 50 μm.

## DISCUSSION

Our semi-systematic review reveals that although there are a small number of studies that either used both sexes or reported data analysis in a sex-aware fashion, there is evidence for sex differences in prostaglandin-mediated effects in both animals and humans. The data suggests that PGE_2_ signaling promotes pain more efficaciously in males, and that COX inhibitors may be more effective in men than in women in experimental pain models. Our experimental data confirms a finding from the rodent literature in humans where we observed that the expression of PTGDS is higher in female human DRG neurons from organ donors. Our findings highlight that there are likely important sex differences in signaling for one of the most widely studied classes of inflammatory mediators, the PGs.

In this review, we analyzed 100 clinical trial studies on the role of PGs and COX inhibitors in pain and inflammation. Most of these studies demonstrated the analgesic efficacy of selective and/or nonselective COX inhibitors. Patient PG levels were often compared with healthy controls, and changes in PG levels were tracked after administering COX inhibitors. Overall, both selective and nonselective COX inhibitors appear to be effective in treating inflammatory pain. However, it is not clear if no sex-differences were found or if the data was simply not analyzed in a sex-aware fashion. In a study of NSAID prescriptions in the Danish population, women received more prescriptions for every NSAID than men [18]. Additionally, a study included in our semi-systematic review found that women report more the use NSAIDs than men [67]. In another study of patients receiving oxycodone, females received higher morphine equivalent daily dose of opioid than men [6]. However, a study on postoperative opioid prescriptions after surgery showed that patient and provider demographic characteristics influenced doses prescribed, with males prescribed higher morphine milligram equivalents [69]. Thus, from the data on prescriptions of analgesics, from NSAIDs to opioids, it is unclear whether women require greater amounts of painkillers to achieve equal relief as men, reflecting sex-differences in drug efficacy, or whether prescribing differences are influenced by social factors.

In addition to clinical pain studies, we analyzed 26 experimental pain studies. While two electrically-induced pain models showed a significant difference in ibuprofen efficacy between men and women [63] [9], no difference was seen in a sunburn model of inflammatory pain [53]. Some studies in both the experimental and clinical pain categories involved participants of only one sex. Interestingly, all the experimental pain studies using only one sex in this analysis happened to include male participants only. While less than a fifth of experimental pain studies analyzed data in a sex-aware fashion, almost no clinical pain studies did so. As mentioned above, the experimental pain studies found sex differences in COX action with a greater effect in men. The single clinical pain study that contained a sex-aware analysis found a difference in pain levels between men and women but did not report any difference in NSAID efficacy. This study on arthroscopic knee surgery observed that the women were at greater risk of developing moderate or severe pain [51]. It is unclear whether the remaining papers omitted any sex-aware analyses because investigators determined there were no differences in the data from male and female participants, or simply because no analyses were attempted in a sex-aware fashion.

In light of the evidence we present here for sex differences in PG action and COX inhibitor efficacy, we suggest that clinical pain studies in this area should analyze and report data in a sex-aware fashion. While most published clinical pain studies do not present data in a sex-aware fashion, these studies often involve a large number of participants allowing for exploratory analyses that can form new hypotheses for further testing. If re-analyzed in a sex-aware fashion, this trove of data could yield significant findings on whether COX inhibitors have a different efficacy between men and women. The findings of our review would lead us to hypothesize greater efficacy in men.

Functional differences in pain and COX-inhibitor efficacy may be at least partially explained by molecular differences in PG pathway receptor and enzyme gene expression. Previous work demonstrates that PTGDS is expressed more highly in female than male mouse DRG, so we evaluated whether this difference is conserved in humans [55]. Our findings show that in human DRG, PTGDS is localized within neurons, and expression is higher in female DRG neurons. Interestingly, among human donors of the same sex, there was some variation in DRG neuron PTGDS expression. Age of the donors, especially in females, as well as disease conditions, could potentially contribute to the variation. In our study, neuronal PTGDS expression in donor #12, a 61-year-old female, was nearly two times lower than that in the five other female donors, whose ages were in the pre-menopausal range. While it is difficult to reach firm conclusions from one donor, the finding suggests that female sex hormones likely regulated PTGDS protein expression. We also show that DP1 protein is highly colocalized in glial cells surrounding neurons in human DRG. This localization suggests that glial cells, likely satellite glial cells, may have an important role in mediating the functions of PGD_2_ and regulating signaling effects by this prostaglandin within the DRG.

Our findings highlight sex differences in prostaglandin effects on pain in both the preclinical and clinical literature that may have implications for development of future pain therapeutics targeting the prostaglandin system. Our work also points to the possibility that commonly used NSAIDs may have less efficacy in women than in men, an effect that may also apply to other pain medications like opioids. Finally, our work validates a sex difference observed in PTGDS expression in rodent DRG in the human DRG. Collectively, this mix of semi-systematic review and experimental data points to shortcomings in our understanding of basic pain mechanisms in women. More emphasis is needed on understanding pain mechanisms in women if we are to adequately treat pain in the population that most frequently suffers from chronic pain disorders.

## Abbreviations

prostaglandins, cyclooxygenases, NSAIDs, sex differences, pain, inflammation

## Acknowledgements

The authors thank the organ donors and their families for their enduring gift. We thank employees of the Southwest Transplant Alliance and members of the Price Lab for coordinating and assisting in DRGs recoveries, respectively.

## Supplementary files

**Suppl. file 1**: **Review and summary of preclinical studies included in semi-systematic review. Tab A** – Studies included in semi-systematic review and the biological sex of the animals/subjects (in chronological order). **Tab B** - Details for studies that analyzed data in a sex-aware fashion. Columns: (1) Paper: Year, first author and PMID of article. (2) Prostaglandins: Specific prostaglandins measured, analyzed, or otherwise mentioned in the paper. (3) Cyclooxygenases: Specific cyclooxygenases measured, analyzed, or otherwise mentioned in the paper. (4) Receptors: Specific prostaglandin receptors studied or otherwise mentioned in the paper. (5) Species: Species of the animal model, including tissues, or human participants, including human tissues, used. (6) Sex: Sex of the study subjects or animals. (7) Number/description: Number of subjects included in the study and characteristics of study subjects or animal models (8) Model: Experimentally induced pain model or clinically occurring pain disease model used. (9) Tissue: Tissues sampled, evaluated, or used in analyses in the study. 10) Injection/Drug administration route: Site of injections or route of administration of drugs. (11) Identified Sex Differences: “No” if sex-aware analysis revealed no mechanistic or functional sex-differences; “Yes” if sex-aware analysis revealed mechanistic and/or functional sex differences.

**Suppl. file 2**: **Number of preclinical studies including male, female or both sexes**.

**Suppl. file 3**: **Review and summary of clinical studies included in semi-systematic review**. Details for all studies included in semi-systematic review (not in chronological order). Columns: (1) Paper: Year, first author and PMID of article. (2) Prostaglandins: Specific prostaglandins measured, analyzed, or otherwise mentioned in the paper. (3) Cyclooxygenases: Specific cyclooxygenases measured, analyzed, or otherwise mentioned in the paper. (4) NSAIDs: Specific NSAIDs measured, analyzed, or otherwise mentioned in the paper. (5) Receptors: Specific prostaglandin receptors studied or otherwise mentioned in the paper. (6) Species: Species of the animal model, including tissues, or human participants, including human tissues, used. (7) Sex: Sex of the study subjects or animals. (8) Number/description: Number of subjects included in the study and characteristics of study subjects or animal models (9) Model: Experimentally induced pain model or clinically occurring pain disease model used. (10) Tissue: Tissues sampled, evaluated, or used in analyses in the study. (11) Injection/Drug administration route: Site of injections or route of administration of drugs. (12) Identified Sex Differences: “N/A” (Not Applicable) if only males, only females, or both males and females without sex-aware analysis were included; “No” if sex-aware analysis revealed no mechanistic or functional sex-differences; “Yes” if sex-aware analysis revealed mechanistic and/or functional sex differences.

**Suppl. file 4**: **Number of clinical studies including male subjects, female subjects or both sexes**.

## REFERENCES

[1] Antonova M, Wienecke T, Olesen J, Ashina M. Pro-inflammatory and vasoconstricting prostanoid PGF2alpha causes no headache in man. Cephalalgia 2011;31(15):1532–1541.

[2] Arout CA, Sofuoglu M, Bastian LA, Rosenheck RA. Gender Differences in the Prevalence of Fibromyalgia and in Concomitant Medical and Psychiatric Disorders: A National Veterans Health Administration Study. J Womens Health (Larchmt) 2018;27(8):1035–1044.

[3] Ashina M, Stallknecht B, Bendtsen L, Pedersen JF, Schifter S, Galbo H, Olesen J. Tender points are not sites of ongoing inflammation -in vivo evidence in patients with chronic tension-type headache. Cephalalgia 2003;23(2):109–116.

[4] Avona A, Burgos-Vega C, Burton MD, Akopian AN, Price TJ, Dussor G. Dural Calcitonin Gene-Related Peptide Produces Female-Specific Responses in Rodent Migraine Models. J Neurosci 2019;39(22):4323–4331.

[5] Avona A, Mason BN, Burgos-Vega C, Hovhannisyan AH, Belugin SN, Mecklenburg J, Goffin V, Wajahat N, Price TJ, Akopian AN, Dussor G. Meningeal CGRP-Prolactin Interaction Evokes Female-Specific Migraine Behavior. Ann Neurol 2021;89(6):1129–1144.

[6] Barrachina J, Margarit C, Muriel J, Lopez-Gil V, Lopez-Gil S, Ballester P, Mira-Lorente L, Agullo L, Peiro AM. Sex Differences in Oxycodone/Naloxone vs. Tapentadol in Chronic Non-Cancer Pain: An Observational Real-World Study. Biomedicines 2022;10(10).

[7] Buisseret B, Alhouayek M, Guillemot-Legris O, Muccioli GG. Endocannabinoid and Prostanoid Crosstalk in Pain. Trends Mol Med 2019;25(10):882–896.

[8] Burke JG, Rw GW, Conhyea D, McCormack D, Dowling FE, Walsh MG, Fitzpatrick JM. Human nucleus pulposis can respond to a pro-inflammatory stimulus. Spine (Phila Pa 1976) 2003;28(24):2685–2693.

[9] Butcher BE, Carmody JJ. Sex differences in analgesic response to ibuprofen are influenced by expectancy: a randomized, crossover, balanced placebo-designed study. Eur J Pain 2012;16(7):1005–1013.

[10] Chen L, Yang G, Grosser T. Prostanoids and inflammatory pain. Prostaglandins Other Lipid Mediat 2013;104-105:58–66.

[11] Chen Z, Huang Q, Song X, Ford NC, Zhang C, Xu Q, Lay M, He S-Q, Dong X, Hanani M. Purinergic signaling between neurons and satellite glial cells of mouse dorsal root ganglia modulates neuronal excitability in vivo. Pain 2022;163(8):1636–1647.

[12] Chillingworth NL, Morham SG, Donaldson LF. Sex differences in inflammation and inflammatory pain in cyclooxygenase-deficient mice. Am J Physiol Regul Integr Comp Physiol 2006;291(2):R327–334.

[13] Cullen L, Kelly L, Connor SO, Fitzgerald DJ. Selective cyclooxygenase-2 inhibition by nimesulide in man. J Pharmacol Exp Ther 1998;287(2):578–582.

[14] Davidson S, Golden JP, Copits BA, Ray PR, Vogt SK, Valtcheva MV, Schmidt RE, Ghetti A, Price TJ, Gereau RWt. Group II mGluRs suppress hyperexcitability in mouse and human nociceptors. Pain 2016;157(9):2081–2088.

[15] Eguchi N, Minami T, Shirafuji N, Kanaoka Y, Tanaka T, Nagata A, Yoshida N, Urade Y, Ito S, Hayaishi O. Lack of tactile pain (allodynia) in lipocalin-type prostaglandin D synthase-deficient mice. Proc Natl Acad Sci U S A 1999;96(2):726–730.

[16] Fillingim RB, King CD, Ribeiro-Dasilva MC, Rahim-Williams B, Riley JL, 3rd. Sex, gender, and pain: a review of recent clinical and experimental findings. J Pain 2009;10(5):447–485.

[17] Fisher JL, Jones EF, Flanary VL, Williams AS, Ramsey EJ, Lasseigne BN. Considerations and challenges for sex-aware drug repurposing. Biol Sex Differ 2022;13(1):13.

[18] Fosbol EL, Gislason GH, Jacobsen S, Abildstrom SZ, Hansen ML, Schramm TK, Folke F, Sorensen R, Rasmussen JN, Kober L, Madsen M, Torp-Pedersen C. The pattern of use of non-steroidal anti-inflammatory drugs (NSAIDs) from 1997 to 2005: a nationwide study on 4.6 million people. Pharmacoepidemiol Drug Saf 2008;17(8):822–833.

[19] Gottesdiener K, Mehlisch DR, Huntington M, Yuan WY, Brown P, Gertz B, Mills S. Efficacy and tolerability of the specific cyclooxygenase-2 inhibitor DFP compared with naproxen sodium in patients with postoperative dental pain. Clin Ther 1999;21(8):1301–1312.

[20] Ito S, Okuda-Ashitaka E, Minami T. Central and peripheral roles of prostaglandins in pain and their interactions with novel neuropeptides nociceptin and nocistatin. Neuroscience research 2001;41(4):299–332.

[21] Ji R-R, Berta T, Nedergaard M. Glia and pain: is chronic pain a gliopathy? Pain® 2013;154:S10–S28.

[22] Langford RM, Mehta V. Selective cyclooxygenase inhibition: its role in pain and anaesthesia. Biomed Pharmacother 2006;60(7):323–328.

[23] Liang X, Wu L, Hand T, Andreasson K. Prostaglandin D2 mediates neuronal protection via the DP1 receptor. J Neurochem 2005;92(3):477–486.

[24] Liu L, Karagoz H, Herneisey M, Zor F, Komatsu T, Loftus S, Janjic BM, Gorantla VS, Janjic JM. Sex Differences Revealed in a Mouse CFA Inflammation Model with Macrophage Targeted Nanotheranostics. Theranostics 2020;10(4):1694–1707.

[25] Minami T, Okuda-Ashitaka E, Mori H, Ito S, Hayaishi O. Prostaglandin D2 inhibits prostaglandin E2-induced allodynia in conscious mice. Journal of Pharmacology and Experimental Therapeutics 1996;278(3):1146–1152.

[26] Minami T, Okuda-Ashitaka E, Nishizawa M, Mori H, Ito S. Inhibition of nociceptin-induced allodynia in conscious mice by prostaglandin D2. Br J Pharmacol 1997;122(4):605–610.

[27] Minami T, Okuda-Ashitaka E, Nishizawa M, Mori H, Ito S. Inhibition of nociceptin-induced allodynia in conscious mice by prostaglandin D2. British journal of pharmacology 1997;122(4):605.

[28] Mogil JS. Laboratory environmental factors and pain behavior: the relevance of unknown unknowns to reproducibility and translation. Lab Anim (NY) 2017;46(4):136–141.

[29] Mogil JS. Qualitative sex differences in pain processing: emerging evidence of a biased literature. Nat Rev Neurosci 2020;21(7):353–365.

[30] Namer B, Schick M, Kleggetveit IP, Orstavik K, Schmidt R, Jorum E, Torebjork E, Handwerker H, Schmelz M. Differential sensitization of silent nociceptors to low pH stimulation by prostaglandin E2 in human volunteers. Eur J Pain 2015;19(2):159–166.

[31] Natura G, Bar KJ, Eitner A, Boettger MK, Richter F, Hensellek S, Ebersberger A, Leuchtweis J, Maruyama T, Hofmann GO, Halbhuber KJ, Schaible HG. Neuronal prostaglandin E2 receptor subtype EP3 mediates antinociception during inflammation. Proc Natl Acad Sci U S A 2013;110(33):13648–13653.

[32] Nishimura M, Segami N, Kaneyama K, Suzuki T, Miyamaru M. Relationships between pain-related mediators and both synovitis and joint pain in patients with internal derangements and osteoarthritis of the temporomandibular joint. Oral Surg Oral Med Oral Pathol Oral Radiol Endod 2002;94(3):328–332.

[33] North RY, Li Y, Ray P, Rhines LD, Tatsui CE, Rao G, Johansson CA, Zhang H, Kim YH, Zhang B, Dussor G, Kim TH, Price TJ, Dougherty PM. Electrophysiological and transcriptomic correlates of neuropathic pain in human dorsal root ganglion neurons. Brain 2019;142(5):1215–1226.

[34] Ogorochi T, Narumiya S, Mizuno N, Yamashita K, Miyazaki H, Hayaishi O. Regional distribution of prostaglandins D2, E2, and F2α and related enzymes in postmortem human brain. Journal of neurochemistry 1984;43(1):71–82.

[35] Ohkubo T, Shibata M, Takahashi H, Inoki R. Effect of prostaglandin D2 on pain and inflammation. Jpn J Pharmacol 1983;33(1):264–266.

[36] Page GG, Ben-Eliyahu S. Indomethacin attenuates the immunosuppressive and tumor-promoting effects of surgery. J Pain 2002;3(4):301–308.

[37] Paige C, Plasencia-Fernandez I, Kume M, Papalampropoulou-Tsiridou M, Lorenzo LE, David ET, He L, Mejia GL, Driskill C, Ferrini F, Feldhaus AL, Garcia-Martinez LF, Akopian AN, De Koninck Y, Dussor G, Price TJ. A Female-Specific Role for Calcitonin Gene-Related Peptide (CGRP) in Rodent Pain Models. J Neurosci 2022;42(10):1930–1944.

[38] Patil M, Hovhannisyan AH, Wangzhou A, Mecklenburg J, Koek W, Goffin V, Grattan D, Boehm U, Dussor G, Price TJ, Akopian AN. Prolactin receptor expression in mouse dorsal root ganglia neuronal subtypes is sex-dependent. J Neuroendocrinol 2019;31(8):e12759.

[39] Paulsen G, Egner IM, Drange M, Langberg H, Benestad HB, Fjeld JG, Hallen J, Raastad T. A COX-2 inhibitor reduces muscle soreness, but does not influence recovery and adaptation after eccentric exercise. Scand J Med Sci Sports 2010;20(1):e195–207.

[40] Racz I, Schutz B, Abo-Salem OM, Zimmer A. Visceral, inflammatory and neuropathic pain in glycine receptor alpha 3-deficient mice. Neuroreport 2005;16(18):2025–2028.

[41] Ray P, Torck A, Quigley L, Wangzhou A, Neiman M, Rao C, Lam T, Kim JY, Kim TH, Zhang MQ, Dussor G, Price TJ. Comparative transcriptome profiling of the human and mouse dorsal root ganglia: an RNA-seq-based resource for pain and sensory neuroscience research. Pain 2018;159(7):1325–1345.

[42] Ray PR, Khan J, Wangzhou A, Tavares-Ferreira D, Akopian AN, Dussor G, Price TJ. Transcriptome Analysis of the Human Tibial Nerve Identifies Sexually Dimorphic Expression of Genes Involved in Pain, Inflammation, and Neuro-Immunity. Front Mol Neurosci 2019;12:37.

[43] Ray PR, Shiers S, Caruso JP, Tavares-Ferreira D, Sankaranarayanan I, Uhelski ML, Li Y, North RY, Tatsui C, Dussor G, Burton MD, Dougherty PM, Price TJ. RNA profiling of human dorsal root ganglia reveals sex-differences in mechanisms promoting neuropathic pain. Brain 2022.

[44] Ricciotti E, FitzGerald GA. Prostaglandins and inflammation. Arterioscler Thromb Vasc Biol 2011;31(5):986–1000.

[45] Rosen S, Ham B, Mogil JS. Sex differences in neuroimmunity and pain. J Neurosci Res 2017;95(1-2):500–508.

[46] Scheff NN, Gold MS. Sex differences in the inflammatory mediator-induced sensitization of dural afferents. J Neurophysiol 2011;106(4):1662–1668.

[47] Sekeroglu A, Jacobsen JM, Jansen-Olesen I, Gupta S, Sheykhzade M, Olesen J, Bhatt DK. Effect of PGD(2) on middle meningeal artery and mRNA expression profile of L-PGD(2) synthase and DP receptors in trigeminovascular system and other pain processing structures in rat brain. Pharmacol Rep 2017;69(1):50–56.

[48] Shaik JS, Miller TM, Graham SH, Manole MD, Poloyac SM. Rapid and simultaneous quantitation of prostanoids by UPLC-MS/MS in rat brain. J Chromatogr B Analyt Technol Biomed Life Sci 2014;945-946:207–216.

[49] Shiers SI, Sankaranarayanan I, Jeevakumar V, Cervantes A, Reese JC, Price TJ. Convergence of peptidergic and non-peptidergic protein markers in the human dorsal root ganglion and spinal dorsal horn. J Comp Neurol 2021;529(10):2771–2788.

[50] Shimizu T, Mizuno N, Amano T, Hayaishi O. Prostaglandin D2, a neuromodulator. Proceedings of the National Academy of Sciences 1979;76(12):6231–6234.

[51] Solheim N, Gregersen I, Halvorsen B, Bjerkeli V, Stubhaug A, Gordh T, Rosseland LA. Randomized controlled trial of intra-articular ketorolac on pain and inflammation after minor arthroscopic knee surgery. Acta Anaesthesiol Scand 2018;62(6):829–838.

[52] Sukoff Rizzo SJ, McTighe S, McKinzie DL. Genetic Background and Sex: Impact on Generalizability of Research Findings in Pharmacology Studies. Handb Exp Pharmacol 2020;257:147–162.

[53] Sycha T, Gustorff B, Lehr S, Tanew A, Eichler HG, Schmetterer L. A simple pain model for the evaluation of analgesic effects of NSAIDs in healthy subjects. Br J Clin Pharmacol 2003;56(2):165–172.

[54] Tavares-Ferreira D, Ray PR, Sankaranarayanan I, Mejia GL, Wangzhou A, Shiers S, Uttarkar R, Megat S, Barragan-Iglesias P, Dussor G. Sex differences in nociceptor translatomes contribute to divergent prostaglandin signaling in male and female mice. Biological Psychiatry 2022;91(1):129–140.

[55] Tavares-Ferreira D, Ray PR, Sankaranarayanan I, Mejia GL, Wangzhou A, Shiers S, Uttarkar R, Megat S, Barragan-Iglesias P, Dussor G, Akopian AN, Price TJ. Sex Differences in Nociceptor Translatomes Contribute to Divergent Prostaglandin Signaling in Male and Female Mice. Biol Psychiatry 2022;91(1):129–140.

[56] Tavares-Ferreira D, Shiers S, Ray PR, Wangzhou A, Jeevakumar V, Sankaranarayanan I, Cervantes AM, Reese JC, Chamessian A, Copits BA, Dougherty PM, Gereau RWt, Burton MD, Dussor G, Price TJ. Spatial transcriptomics of dorsal root ganglia identifies molecular signatures of human nociceptors. Sci Transl Med 2022;14(632):eabj8186.

[57] Theken KN, Hersh EV, Lahens NF, Lee HM, Li X, Granquist EJ, Giannakopoulos HE, Levin LM, Secreto SA, Grant GR, Detre JA, FitzGerald GA, Grosser T, Farrar JT. Variability in the Analgesic Response to Ibuprofen Is Associated With Cyclooxygenase Activation in Inflammatory Pain. Clin Pharmacol Ther 2019;106(3):632–641.

[58] Uda R, Horiguchi S, Ito S, Hyodo M, Hayaishi O. Nociceptive effects induced by intrathecal administration of prostaglandin D2, E2, or F2 alpha to conscious mice. Brain Res 1990;510(1):26–32.

[59] Urade Y, Hayaishi O. Prostaglandin D2 and sleep regulation. Biochimica et Biophysica Acta (BBA)-Molecular and Cell Biology of Lipids 1999;1436(3):606–615.

[60] Vetvik KG, MacGregor EA. Sex differences in the epidemiology, clinical features, and pathophysiology of migraine. Lancet Neurol 2017;16(1):76–87.

[61] Vina ER, Kwoh CK. Epidemiology of osteoarthritis: literature update. Curr Opin Rheumatol 2018;30(2):160–167.

[62] Vonkeman HE, van de Laar MA. Nonsteroidal anti-inflammatory drugs: adverse effects and their prevention. Semin Arthritis Rheum 2010;39(4):294–312.

[63] Walker JS, Carmody JJ. Experimental pain in healthy human subjects: gender differences in nociception and in response to ibuprofen. Anesth Analg 1998;86(6):1257–1262.

[64] Wang TA, Teo CF, Åkerblom M, Chen C, Tynan-La Fontaine M, Greiner VJ, Diaz A, McManus MT, Jan YN, Jan LY. Thermoregulation via temperature-dependent PGD2 production in mouse preoptic area. Neuron 2019;103(2):309-322. e307.

[65] Wangzhou A, McIlvried LA, Paige C, Barragan-Iglesias P, Shiers S, Ahmad A, Guzman CA, Dussor G, Ray PR, Gereau RWt, Price TJ. Pharmacological target-focused transcriptomic analysis of native vs cultured human and mouse dorsal root ganglia. Pain 2020;161(7):1497–1517.

[66] Wangzhou A, Paige C, Neerukonda SV, Naik DK, Kume M, David ET, Dussor G, Ray PR, Price TJ. A ligand-receptor interactome platform for discovery of pain mechanisms and therapeutic targets. Sci Signal 2021;14(674).

[67] Wilcox CM, Cryer B, Triadafilopoulos G. Patterns of use and public perception of over-the-counter pain relievers: focus on nonsteroidal antiinflammatory drugs. The Journal of rheumatology 2005;32(11):2218–2224.

[68] Yuan Q, Liu X, Xian Y-f, Yao M, Zhang X, Huang P, Wu W, Lin Z-X. Satellite glia activation in dorsal root ganglion contributes to mechanical allodynia after selective motor fiber injury in adult rats. Biomedicine & pharmacotherapy 2020;127:110187.

[69] Zaveri S, Nobel TB, Khetan P, Srinivasan M, Divino CM. Surgeon Bias in Postoperative Opioid Prescribing. World J Surg 2022;46(7):1660–1666.

[70] Zeisel A, Hochgerner H, Lonnerberg P, Johnsson A, Memic F, van der Zwan J, Haring M, Braun E, Borm LE, La Manno G, Codeluppi S, Furlan A, Lee K, Skene N, Harris KD, Hjerling-Leffler J, Arenas E, Ernfors P, Marklund U, Linnarsson S. Molecular Architecture of the Mouse Nervous System. Cell 2018;174(4):999–1014 e1022.

[71] Zhang P, Bi RY, Gan YH. Glial interleukin-1beta upregulates neuronal sodium channel 1.7 in trigeminal ganglion contributing to temporomandibular joint inflammatory hypernociception in rats. J Neuroinflammation 2018;15(1):117.

